# Climate-driven warming disrupts the symbiosis of bobtail squid *Euprymna scolopes* and the luminous bacterium *Vibrio fischeri*

**DOI:** 10.1101/2025.02.14.638271

**Authors:** E. Otjacques, B. Jatico, T.A. Marques, J.C. Xavier, E. Ruby, M. McFall-Ngai, R. Rosa

**Affiliations:** MARE – Marine and Environmental Sciences Centre/ARNET - Aquatic Research Network, Laboratório Marítimo da Guia, Faculdade de Ciências, Universidade de Lisboa, Cascais, Portugal; Division of Biology & Biological Engineering, Carnegie Science, California Institute of Technology, Pasadena CA, United States of America; MARE – Marine and Environmental Sciences Centre/ARNET – Aquatic Research Network, Department of Life Sciences, Universidade de Coimbra, Coimbra, Portugal; Departamento de Biologia Animal, Faculdade de Ciências, Universidade de Lisboa, Lisboa, Portugal; Centro de Estatística e Aplicações, Faculdade de Ciências, Universidade de Lisboa, Lisboa, Portugal; Center for Research into Environmental Ecological Modelling, University of St Andrews, St Andrews, Scotland; British Antarctic Survey, Natural Environment Research Council, High Cross, Madingley Road, Cambridge, UK

**Keywords:** Sepiolidae, Cephalopod, Symbiosis, Temperature, Bioluminescence

## Abstract

Under the current climate crisis, marine heatwaves (MHW) are expected to intensify and become more frequent in the future, leading to adverse effects on marine life. Here, we aimed to investigate the impact of environmental warming on the symbiotic relationship between the Hawaiian bobtail squid (*Euprymna scolopes*) and the bioluminescent bacterium *Vibrio fischeri*. We exposed eggs of *E. scolopes* to three different temperatures during the embryogenesis, namely: i) 25 °C (yearly average), ii) 27 °C (average summer), or iii) 30 °C (category IV MHW), followed by a colonisation assay under the same conditions. Decreased hatching success and reduced developmental time were observed across warmer conditions compared to 25 °C. Moreover, exposure to the category IV MHW led to a significant decrease in survival after 48 hours. With increasing temperature, bobtail squids required more bacteria in the surrounding seawater for successful colonisation. When colonised, the regression of the light organ’s appendages was not dependent on temperature, but the opposite was found in non-colonised bobtail squids. Furthermore, the capacity to maintain the symbiosis diminished significantly with temperature. Finally, the capacity for crypt 3 formation in the squid’s light organ, which is crucial for enhancing resilience under stress, also declined with warming conditions. This study emphasises the critical need to study the dynamics of microbial symbiosis under the projected conditions for the ocean of tomorrow.

## 1 Introduction

Eukaryotes are known to form symbiotic associations with bacteria, which can be deleterious (i.e., parasitism) or beneficial to one or both partners (i.e., bacterial commensalism or mutualism, respectively (Sachs, Skophammer, et al., 2011)). Beneficial symbiosis depends on one of two main modes of transmission: i) vertical transmission involves the transfer of the symbiont from the host to its offspring during embryogenesis, or ii) horizontal transmission refers to the host obtaining the symbiont from the environment (Sachs, Essenberg, et al., 2011; Sachs, Skophammer, et al., 2011; Russell, 2019). Horizontally transmitted symbioses (HTS) are usually initiated soon after embryogenesis and birth (Koenig et al., 2011; Essock-Burns et al., 2023). However, such an association can occur at a specific life stage of the host, following a short or extended timeframe for colonisation, or throughout the host’s lifetime (Bright & Bulgheresi, 2010). As parasites and other deleterious microbes can be treated and influenced by environmental factors, microorganisms can also be affected (Shehata et al., 2013; Allen, 2017; Xia et al., 2022). Therefore, changes in the microbiome regulation may lead to negative outcomes, such as inflammatory responses and a general degradation of the host’s health (Iebba et al., 2012; Rosenfeld, 2017; Weiss & Hennet, 2017; Sugihara & Kamada, 2024).

Extreme events, such as marine heatwaves (MHW), are expected to increase in intensity and to become more frequent in the future (Frölicher et al., 2018; Oliver et al., 2021). Tropical and subtropical regions are, in fact, expected to endure the strongest changes (Frölicher et al., 2018). MHW are events that occur alongside global ocean warming, defined as a prolonged period of abnormally warm water (Frölicher et al., 2018; Hobday et al., 2018; Oliver et al., 2021). In addition to socioeconomic impacts, MHW are a major threat to marine ecosystems, affecting species at an individual level (e.g., increased mortality) and provoking shifts in population and a global loss of biodiversity (Smale et al., 2019; Smith et al., 2023). MHW are known to cause mass bleaching events amongst corals and sponges through the expulsion of the symbionts from the host (Bell et al., 2023; Smith et al., 2023). Moreover, sponges and octocorals have been shown to undergo a reorganisation of their microbiome in response to MHW by expelling certain bacteria over others (Prioux et al., 2023; Bell et al., 2024). In fact, octocorals exhibit an increase in pathogenic bacteria (Prioux et al., 2023), suggesting an increasing vulnerability of symbiotic organisms to such events. Yet, it has also been shown that symbiotic endolithic microbes reduce mussel vulnerability to MHW conditions (Zardi et al., 2024) and that the vertical transmission of microbes from heatwave-exposed organisms may provide resilience for following generations (Baldassarre et al., 2022; Strano et al., 2023).

Cephalopods play important economic and ecological roles in marine ecosystems, with a central status in trophic food webs as both predators and preys (Boyle & Rodhouse, 2005; Arkhipkin et al., 2015; de la Chesnais et al., 2019; González & Pierce, 2021; Sauer et al., 2021; Bobowski et al., 2023). Like other marine ectothermic species, cephalopods are affected by environmental warming (Borges et al., 2023), especially during early ontogeny (Pimentel et al., 2012; Repolho et al., 2014; Rosa et al., 2012, 2014). Increased temperature generally reduces developmental time (i.e., the number of days until hatching) and survival of early life stages (Pimentel et al., 2012; Rosa et al., 2013; Repolho et al., 2014; Borges et al., 2023). Although we have extended knowledge of climate change-driven changes in cephalopods (Borges et al., 2023), the impact of MHW on cephalopods living in symbiosis (e.g., associated with bioluminescent bacteria, ∼ 15 % of cephalopods (Otjacques et al., 2023)) is unknown.

In this study, we used the Hawaiian bobtail squid *Euprymna scolopes* to investigate the establishment of bacterial symbiosis under MHW conditions. This sepiolid is typically found in sandy and extremely shallow coastal waters, although the actual range of this species is difficult to assess as it can inhabit areas extending to the continental shelf (∼ 200 m (Berry, 1914; Anderson & Mather, 1996; Anderson et al., 2000; Stabb & Millikan, 2008)). It is known for its binary association with *Vibrio fischeri*, a bioluminescent bacterium (Nyholm & McFall-Ngai, 2021; Visick et al., 2021). During the embryogenesis, the duration of which is temperature-dependent (Lee et al., 2009), *E. scolopes* develops a bilobed light organ (LO) composed of six crypt spaces (three crypts on each side of the LO, named C1, C2 and C3. These crypts are the final location for the establishment of the bacteria. They are formed gradually during the embryogenesis by invagination of the epithelial surface (Montgomery & McFall-Ngai, 1993). The formation of C3 begins only a few days before hatching (Montgomery & McFall-Ngai, 1993). Upon hatching, the bobtail squid will acquire the bacterial symbiont horizontally from the surrounding environment, followed by a profound morphogenesis of the LO (e.g., regression of ciliated appendages) within 96 hours (Montgomery & McFall-Ngai, 1994; Nyholm & McFall-Ngai, 2004). Once the symbiosis is well-established, the animal uses the bacterial bioluminescence at night for counterillumination, hiding its silhouette as a form of camouflage (Jones & Nishiguchi, 2004). Although the symbiosis between *E. scolopes* and *V. fischeri* is well understood, there is a lack of knowledge on the impact of environmental stressors, such as temperature, on the initiation of this symbiotic association. Moreover, because it is suggested that the colonisation of C3 offers resilience to the host, allowing the symbiosis to be recovered after disturbances (Essock-Burns et al., 2023), it is important to understand the influence of environmental changes (i.e., temperature) on the crypt formation, potentially affecting the resilience of this species.

Here, we aim to investigate the resilience of the symbiotic relationship between *E. scolopes* and *Vibrio fischeri* to different temperature conditions (i.e., winter temperature, summer temperature and a category IV marine heatwave) during embryogenesis. We hypothesise: i) lower hatching success associated with increased temperature, as well as a decrease in developmental time with warmer conditions, ii) once hatching, a decreased survival rate over time with increased temperature, iii) decreased colonisation efficiency and effective colonisation under MHW conditions, and iv) that light organ morphology and crypts formation will be affected with increasing temperature.

## 2 Material and methods

### 2.1 Marine heatwave modelling

The model for marine heatwaves was built using a 31-year dataset (1993-2024) of sea surface temperatures (SSTs) collected from the Copernicus Marine Service (21.25 °N, 157.75 °W). The R package “heatwaveR” v.0.4.6 (Schlegel & Smit, 2018) was used to create the temperature curves assessing yearly temperature variation and temperatures for each level of marine heatwaves (MHW). This package follows the definition offered by Hobday et al. (2018) with which a category IV MHW occurs when SSTs exceed the expected temperature by four times (Supplementary Figure 1).

### 2.2 Experimental design and set-up

A breeding colony of Hawaiian bobtail squid was maintained in artificial seawater (Instant Ocean) at the California Institute of Technology (Pasadena, CA, USA). The adults were obtained in February 2024 from Paikō Bay on the southern coast of O’ahu, Hawai’i (21.281 °N, 157.731 ° W). A total of three egg clutches, from different adult females were used during this experiment in March 2024. For each clutch, an approximate equal number of eggs was placed in nine separate mesh bags. One bag per clutch, three bags in total, were placed in 18-L containers (three containers per temperature, nine containers in total). The containers were equipped with either a 50 W or 100 W water heater, depending on the final temperature condition, with an external temperature controller (accuracy ± 0.55 °C) and an air stone to ensure water oxygenation. The photoperiod was kept under a 12-h light:12-h dark cycle, as observed in O’ahu for March. Temperature (± 0.1°C, Apera Instruments), pH (± 0.01 unit, Apera Instruments), dissolved oxygen (± 0.01 mg/L or ± 0.1 %, Apera Instruments) and salinity (± 1 PSU, NISupply) were measured daily (Supplementary Table 1). To ensure good water quality, we replaced 6 L of the seawater every two days. However, to allow natural hatching of the bobtail squid without the disturbance that could be caused from water changes, these were interrupted as soon as the first animal hatched in each container.

The eggs were reared until hatching at one of the following temperatures: i) 25 °C; ii) 27 °C; iii) 30 °C (Supplementary Table 1). The temperatures were selected based on the previously described marine heatwave model, for which 25 °C and 27 °C correspond approximatively to the yearly average temperature and the maximum temperature in a year, respectively. 30 °C corresponds approximatively to the highest temperature during a category IV MHW (Supplementary Figure 1). Temperature was increased with an increment of 1 °C per day until reaching the final temperature of each treatment, then maintained constant until the end of the experiment (Supplementary Figure 2).

### 2.3 Bobtail squid colonisation assay

To visualise the bacteria under the confocal microscopy, we performed all colonisation with the strain *Vibrio fischeri* ES114 pVSV102, presenting the plasmid carrying a green fluorescent protein (GFP) marker. A single colony was cultured at room temperature overnight in Luria-Bertani salt medium (LBS), supplemented with kanamycin antibiotic at a concentration of 50 µg/mL to maintain the selection pressure of the strain. The overnight culture was sub-cultured in seawater-tryptone medium (SWT) and grown at 28 °C with 220 rpm shaking, until reaching an optical density at 600 nm (OD_600_) of ∼ 0.7. For the colonisation assay, the SWT culture was diluted to reach a final OD_600_ of ∼ 0.2.

To understand the colonisation efficiency at the different temperatures, daily hatchlings were individually inoculated with a three-hour exposure to the GFP-labelled ES114, in 5 mL of natural offshore seawater. Two fresh inocula were prepared daily, with a 10-fold difference, ranging from 235 to 1.2 x 10^4^ Colony-Forming-Unit mL^-1^ (CFU mL^-1^). After three hours, the animals were washed twice by transferring them to fresh artificial seawater (AS) before a final transfer into 5 mL fresh AS. To monitor colonisation status, the animal’s bioluminescence was measured using a TD 20/20 luminometer (Turner Designs, Inc.) at five timepoints (i.e., 18-, 24-, 30-, 42- and 48-h post inoculation [hpi]). The animals were transferred into fresh AS after each timepoint. At 48 hpi, we sampled the animals to either i) analyse the light organ (LO) morphology using confocal microscopy or ii) assess the CFU per light organ (Supplementary Figure 2). In the former case, some animals were anesthetised in seawater containing 2% ethanol followed by an overnight fixation in 4% paraformaldehyde (PFA) in marine phosphate-buffered saline (mPBS: 450 mM NaCl, 50 mM sodium phosphate buffer [pH 7.4]). Following the overnight fixation, the fixed samples were washed three times in mPBS for 30 min before storage at 4 °C, until further processing. We dissected the LO from all fixed animals before analysis. In the latter, the other subset of animals was flash-frozen and stored at - 80 °C. To assess the presence of bacteria in each sample, the frozen samples were thawed and homogenised in 500 µL of SWT. Dilutions of the homogenate were spread onto LBS agar medium, and the CFU per light organ were calculated.

### 2.4 Sample preparation and microscopy

The LO were permeabilised and stained in 0.1% Triton X-100 in mPBS in the dark at 4°C on a shaker for two days. We used TO-PRO3 Iodide (ThermoFisher Scientific) to stain the nuclei (dilution 1:1000, excitation/emission [Ex/Em] = 642/661 nm) and Rhodamine 405 (ThermoFisher Scientific) to stain F-actin (dilution 1:40, Ex/Em = 405/450 nm). Stained samples were mounted in Vectashield (Vector Laboratories) and overlaid with a coverslip number 1.5 (Fisher Scientific). Laser scanning confocal microscopy was performed using a Zeiss LSM 900 microscope (Plan-Apochromat 10x/0.45 Air M27 objective).

Confocal z-stacks were analysed using Fiji (ImageJ). LO appendages’ length and area were measured manually (from the edge of the pore to the tip of the appendage; Supplementary Figure 3) using the ‘Freehand’ lines and selection tools, respectively. The measurements were determined using the ‘measure’ function, calibrated to the µm/pixel scale of the original z-stack. The presence of GFP-labelled bacteria was investigated as a sign of colonisation. Finally, the formation status of crypt 3 (C3) was determined for each sample.

### 2.5 Colonisation efficiency assessment

The animals were considered colonised if one of the four conditions was met: i) the luminescence readings ≥ 10 relative-light-unit (RLU) for at least one time point, ii) the number of cells per light organ ≥ 100 CFU, iii) GFP-labelled bacteria were visible on the confocal images, or iv) the LO appendages’ regression was induced (i.e., the appendage length ≤ cut-off value per treatment). This value was measured for each temperature as the appendage length’s standard deviation subtracted from the median value (Supplementary Table 2). Moreover, the colonisation was considered effective when the luminescence readings ≥ 10 RLU at least one time point.

When applicable, we assessed the maintenance of the symbiosis. The colonisation was considered maintained, at 48 hpi specifically, when visible bacteria was observed either by CFU per LO ≥ 100, or by the presence of GFP-labelled bacteria. Additionally, when RLU ≥ 10 at 48 hpi specifically, the symbiosis was also considered maintained, as any loss of the symbiont at this final timepoint may have occurred due to stress-induced sampling of the animal. In case the animal was considered colonised, but no bacteria were observed at 48 hpi, the symbiosis was considered lost (Supplementary Table 3).

### 2.6 Data analysis and statistics

Hatching success and developmental time were analysed on all animals. Because bobtail squids principally hatch around dusk and colonise in the following hours (Nyholm & McFall-Ngai, 2004), we have used animals that hatched five hours pre-to three hours post-dusk in all other analyses (i.e., survival, regression of appendages, colonisation, maintenance of the symbiosis, formation of crypt 3).

Time-to-event data, namely hatching success and survival, were treated using a survival analysis. Kaplan–Meier plots were made to illustrate the survival curves, using the function “ggsurvplot” from the R package “survminer” v. 0.4.9 (Kassambara et al., 2016). Survival curves were visually assessed for violation of the assumption of proportional hazard rates, demonstrated by crossing of the survival curves. The visual crossing of survival curves was confirmed using the function “crosspoint” from the R package “ComparisonSurv” v. 1.1.1 (Lyu et al., 2019). Because of the curves crossing, the two-stage test was used to analyse the hatching success, through pairwise analyses using the function “twostage” from the R package “TSHRC” v. 0.1-6 (Sheng et al., 2008). On the other hand, since there was no visible crossing when assessing the survival of bobtail squids, we used the R package “survival” v. 3.6-4 (Therneau, 2024) and the Cox proportional hazards regression model using the function “coxph”. We used the temperature treatment as a categorical fixed effect for the proportional hazards Cox mixed effects model fit by maximum likelihood, and female was set as a random factor. We verified compliance with the assumption of proportional hazards using the global test statistic in the function “coxph” and graphically using a smoothed spline plot of the Schoenfeld residuals relative to time (Supplementary Figure 4). Finally, post-hoc multiple comparisons were performed using the LogRank test, and p-values were adjusted through Bonferroni–Hochberg corrections to avoid type I errors.

The colonisation efficiency, effective colonisation, the maintenance of the symbiosis and the formation of C3 were analysed using general linear mixed models (GLMM) with a binomial distribution. Colonisation efficiency (i.e., presence of symbiont) and effective colonisation (i.e., presence of bioluminescence) used temperature treatment as the continuous factor, the inoculum size as a continuous covariate, and female as a random factor. The maintenance of the symbiosis was analysed with colonised bobtail squids only, using temperature treatment as a continuous fixed factor and female as a random factor. Finally, the formation of C3 was analysed using temperature treatment as the continuous factor, the developmental time as a covariate and female as a random factor (a total of 7 samples were removed from the dataset, as the detection of C3 could not be made with certainty). We used the function “glmmTMB” from the R package of the same name (v. 1.1.9, (Brooks et al., 2017)) to create the statistical models.

The regression of the light organ appendages was analysed using linear mixed-effects models (LMM, considering a Gaussian response). We used the temperature treatment as a continuous factor, the colonisation status as a covariate and female as a random factor (a total of 19 samples were removed from the dataset as the length could not be measured on the two sides of the light organ). We used the function “lmer” from the R package “lme4” v. 1.1-35.3 (Bates et al., 2015) to fit the model. All models were tested for statistical significance using the function “Anova” from the package “car” v. 3.1-2 (Fox & Weisberg, 2019). All statistical analyses and graphs were performed in R, version 4.3.3 (R Core Team, 2024).

## 3 Results

### 3.1 Hatching success

Warmer temperatures elicited a significant decline in hatching success (Figure 1; Supplementary Table 4.A-B). While 94.2 % of bobtail squids hatched at 25 °C (n_hatching_25_/n_total_25_ = 162/172), there was a significant decrease (i.e., 85.5 %) of hatched animals at 27 °C (n_hatching_27_/n_total_27_ = 147/172, p-value < 0.001), and only 67.8 % hatched when the embryogenesis occurred at 30°C (n_hatching_30_/n_total_30_ = 116/171, p-value = 0.025). Bobtail squids raised at 30 °C also presented a significantly lower hatching success than at 27 °C (p-value = 0.027).

**Figure 1.**
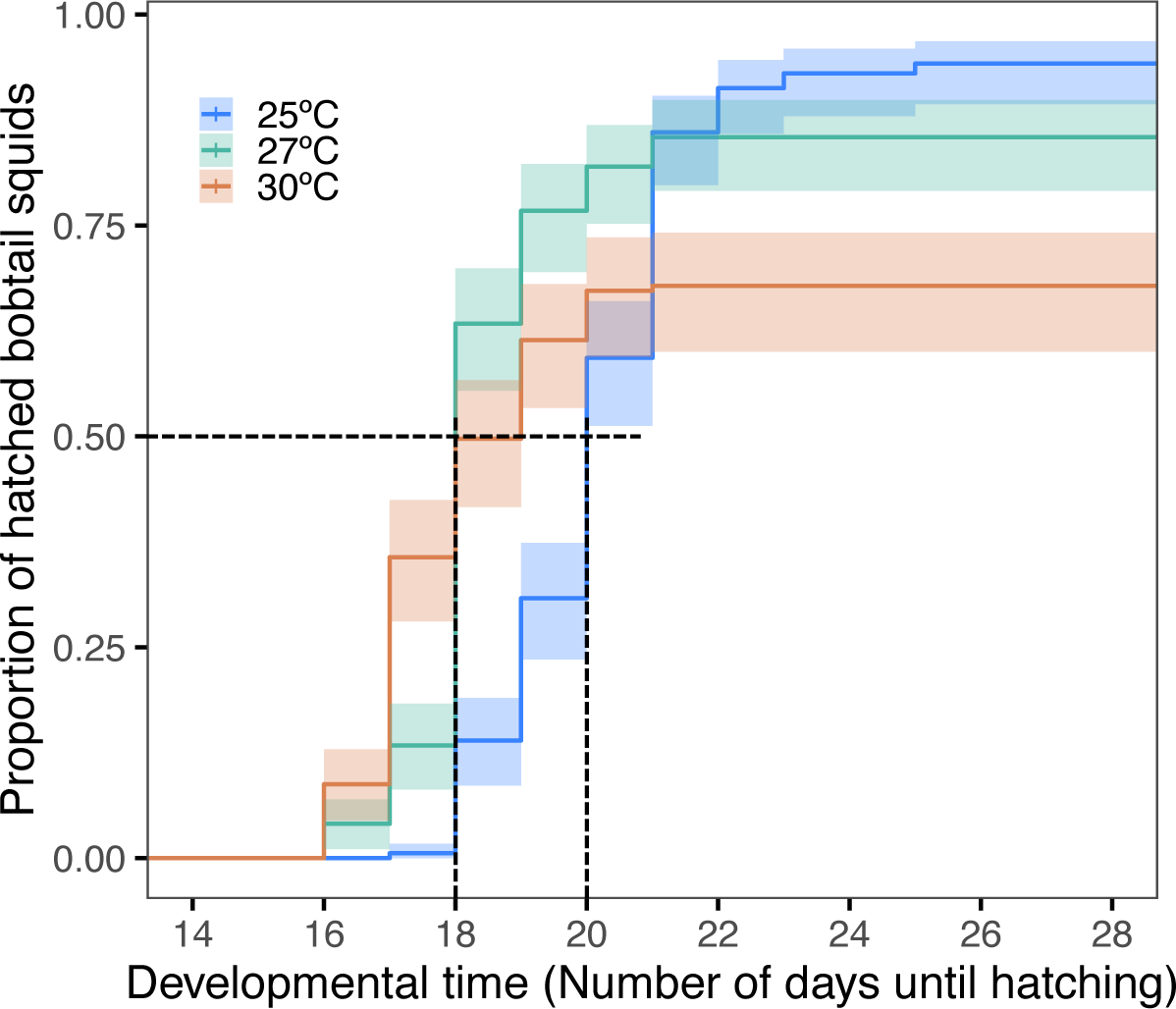
Hatching success of Hawaiian bobtail squid as a function of temperature, measured according to the number of days until hatching (developmental time). The presence of hatchlings was verified daily. The Kaplan-Meier curves show the survival trajectories according to each temperature (blue = 25 °C, green = 27 °C, red = 30 °C). The lines represent the additive proportion of hatched bobtail squid on each day. The shaded area illustrates the 95 % confidence intervals. The dotted lines indicate the developmental time to reach 50 % of hatched bobtail squids, based on 100 % hatching success.

Moreover, we observed a reduction in developmental time (i.e., the number of days until hatching) when exposed to increased temperature (Supplemental Table 5). The median developmental time was 20 ± 1 days when the animals were reared at 25 °C. However, bobtail squids raised at 27 °C showed a median developmental time reduced to 18 ± 1 days (Figure 1). Finally, because of the reduced hatching success due to an embryogenesis at 30 °C, the animals reared under these conditions had a median developmental time of 17 ± 1 days.

### 3.2 Survival

We observed a significant decrease in survival with temperature after hatching (Figure 2; Supplementary Table 6.A-B), throughout 48 h, the proportion of bobtail squids that survived was only 48.9 when exposed to 30 °C (n_survivor_30_/n_hatched_30_ = 44/90), a significantly lower proportion compared to that at 25°C (98.4 %, n_survivor_25_/n_hatched_25_ = 122/124, p-value < 0.001) or 27 °C (94.2 %, n_survivor_27_/n_hatched_27_ = 113/120, p-value < 0.001). No significant difference in survival was observed between the treatment 25 °C and 27 °C (p-value = 0.081, Supplementary Table 6.C).

**Figure 2.**
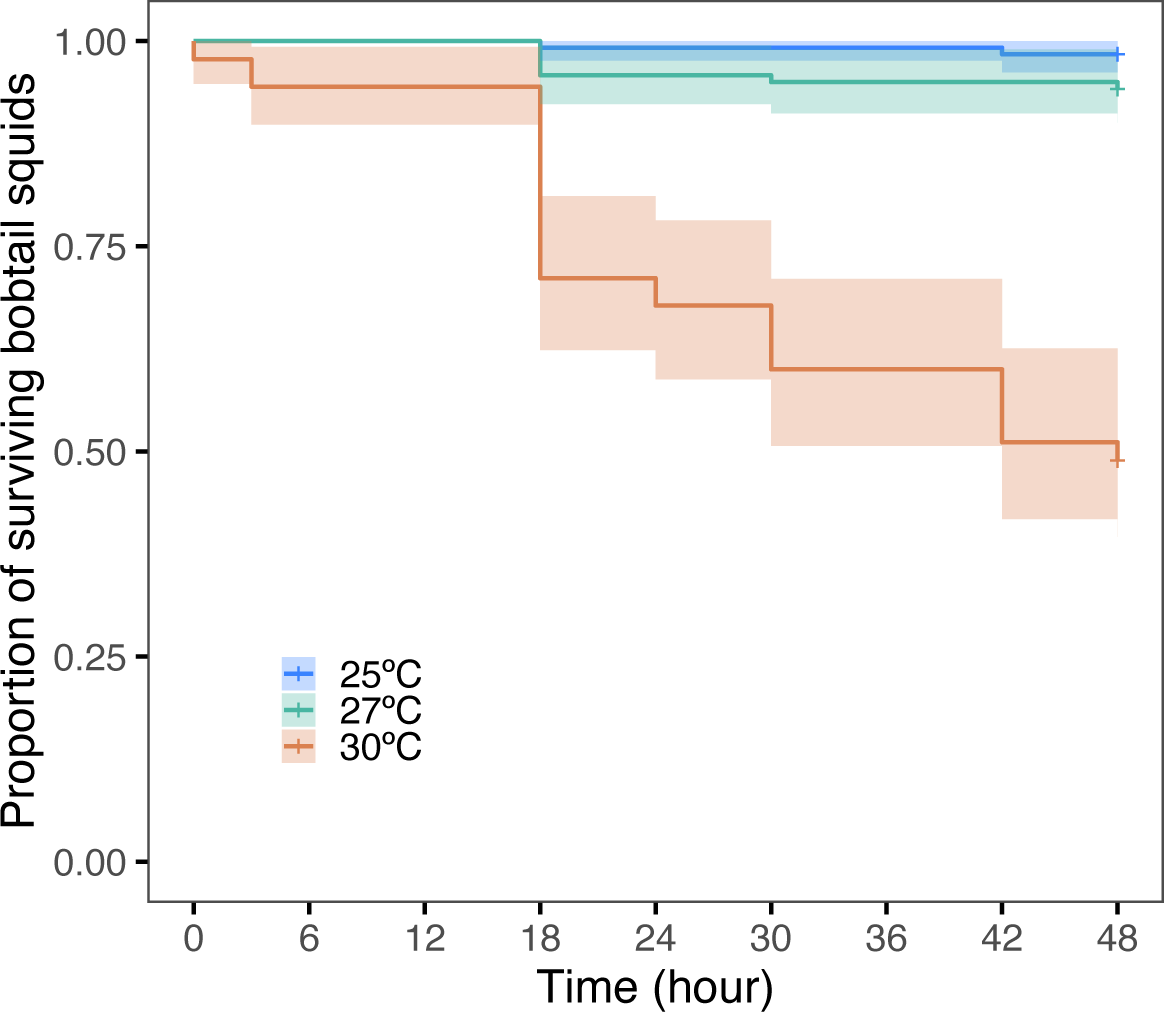
Survival rate of Hawaiian bobtail squid as a function of temperature, measured over the course of 48 h. Mortality was measured at 0 h (death at arrival), 3 h post-inoculation (hpi, measured after the start of colonisation assay), 18, 24, 30, 42 and 48 hpi. The Kaplan-Meier survival trajectories show the survival trajectories according to each temperature (blue = 25 °C, green = 27 °C, red = 30 °C). The lines represent the survival rate on each day. The shaded area illustrates the 95 % confidence intervals.

### 3.1 Colonisation efficiency and appendage regression

Both measures of colonisation efficiency and effective colonisation were statistically significant based on the combined effects of inoculum size and temperature (p-value = 0.022 and 0.001, respectively; Supplementary Table 7.A-B). These results indicate that bobtail squid specimens require an increased number of bacterial cells (i.e., a higher inoculum) in the seawater under warming conditions to achieve successful colonisation (Figure 3.A). These results were similar for effective colonisation (i.e., when bioluminescence is produced; Figure 3.B).

**Figure 3.**
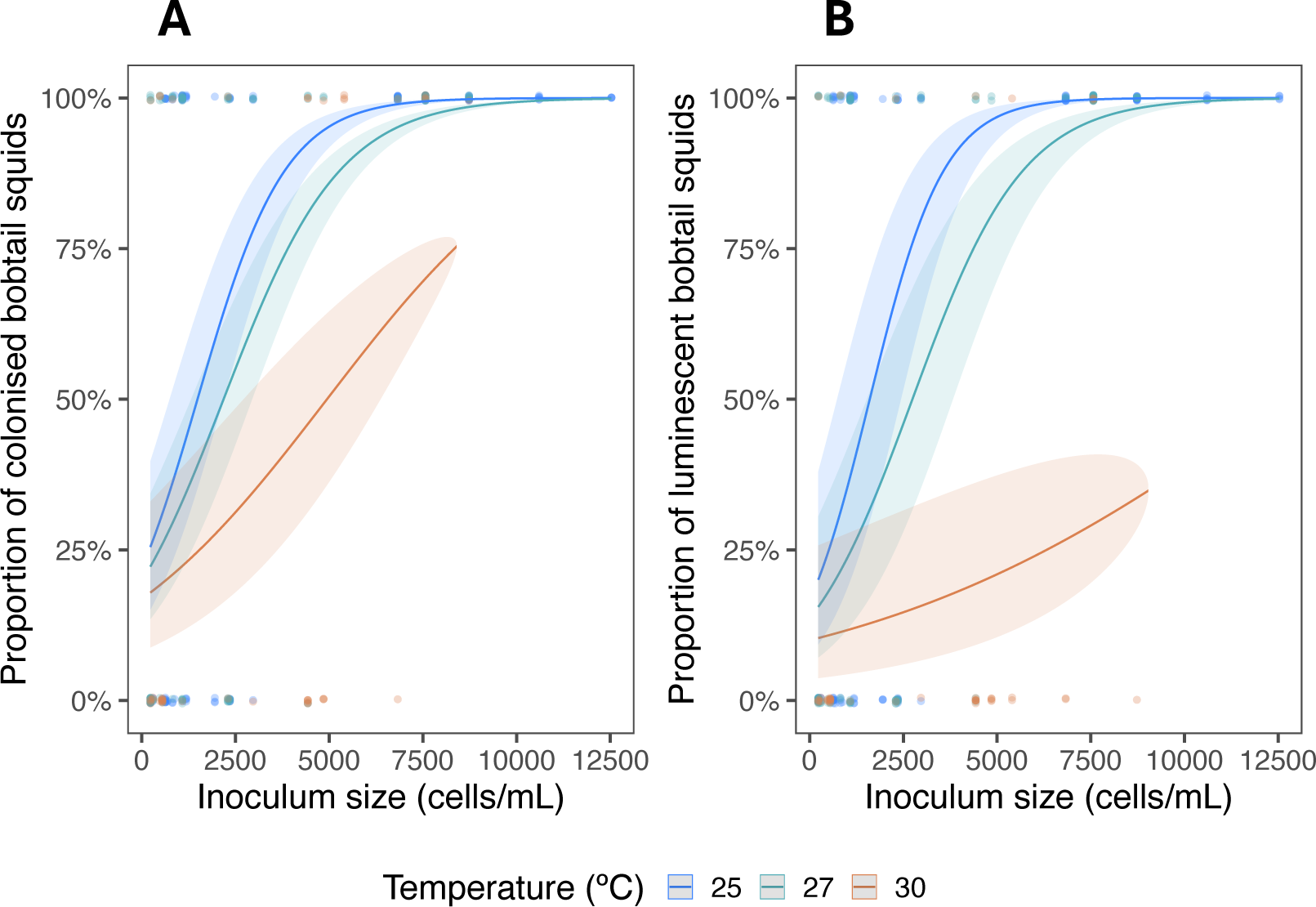
Colonisation efficiency (A) and effective colonisation (B) at different inoculum sizes, according to temperature. Colonisation of bobtail squid specimens by a GFP-labelled ES114 strain. The animals were individually incubated for three h with different concentrations of bacteria (cells/mL) in offshore seawater and transferred into uninoculated offshore seawater for 48 h. The relative light unit (RLU, measure of luminescence) of each animal was measured at 18 h post-inoculation (hpi), 24 hpi, 30 hpi, 42 hpi and 48 hpi. (A) Bobtail squid specimens were considered colonised if one of the four conditions was met in at least one time point: i) the luminescence readings ≥ 10 relative-light-unit (RLU), ii) the number of cells per light organ ≥ 100 CFU, iii) GFP-labelled bacteria were visible in the confocal images, or iv) the LO appendage regression was induced, where the appendage length ≤ cut-off value per treatment. This value was measured for each temperature as the standard deviation of the appendage length subtracted from the median value. (B) The colonisation was considered effective (i.e., the animal was luminescent) when RLU > 10, in at least one of the timepoints mentioned above. Back-transformed predicted means ± 95 % confidence interval from the model and raw data values are presented.

Moreover, the light organ appendage length varied as a function of the combined effects of temperature and colonisation conditions (p-value < 0.001). There was a significant decrease in the appendage length with increasing temperature when the bobtail squid specimens were not colonised. However, temperature did not affect the appendage length when the bobtail squid specimens were colonised (Figure 4, Supplementary Table 8.A). Similar results were obtained for the area of the light organ appendages (Supplementary Figure 5, Supplementary Table 8.B).

**Figure 4.**
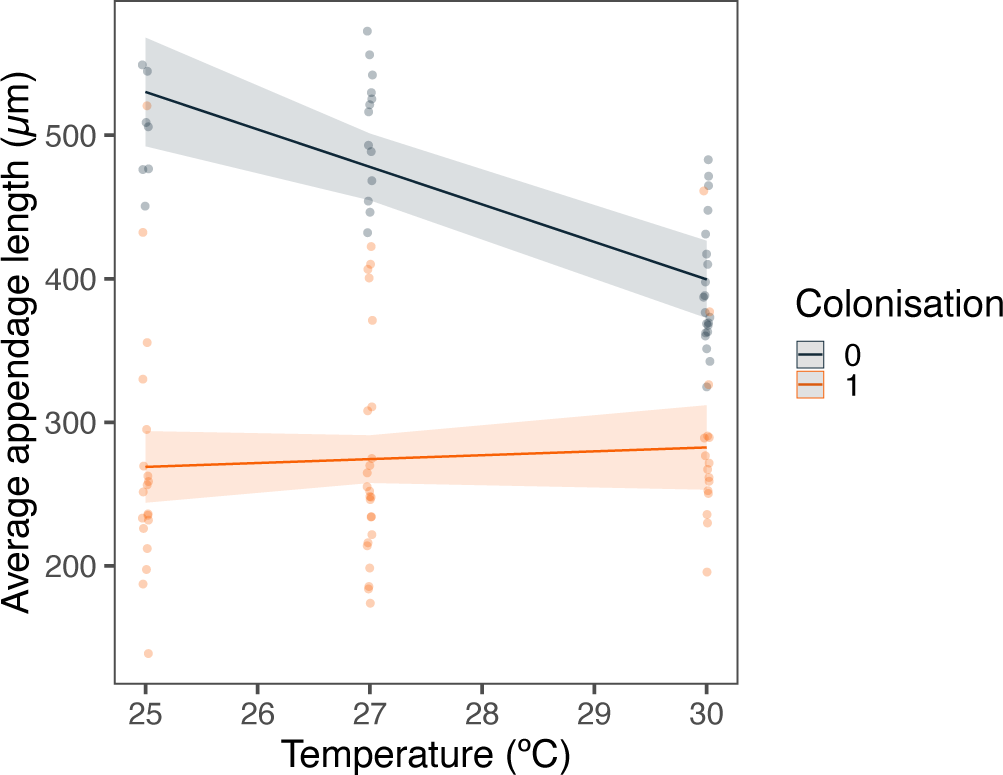
Light-organ appendages length according to temperature and colonisation. The morphology of the light organ, the length of the appendages more specifically, was analysed through a linear mixed model from the Gaussian family, as a function of the temperature and the colonisation. Non-colonised (0) bobtail squids are in black, and colonised (1) bobtail squids are in red. Back-transformed predicted means ± 95% confidence interval from the model and raw data values are presented. The lengths of both appendages were averaged for each animal.

### 3.2 Maintenance of the symbiosis

At 48 hpi, we observed a significant decrease in the maintenance of the symbiosis with increasing temperature (Figure 5; Supplementary Table 9.A-B). 93.6 % of symbiotic animals maintained the symbiosis at 25 °C (n_maintenance_25_/n_total_25_ = 73/78), but it decreased to 82.9 % at 27 °C (n_maintenance_27_/n_total_27_ = 58/70), and to only 29.4 % at 30 °C (n_maintenance_30_/n_total_30_ = 5/17).

**Figure 5.**
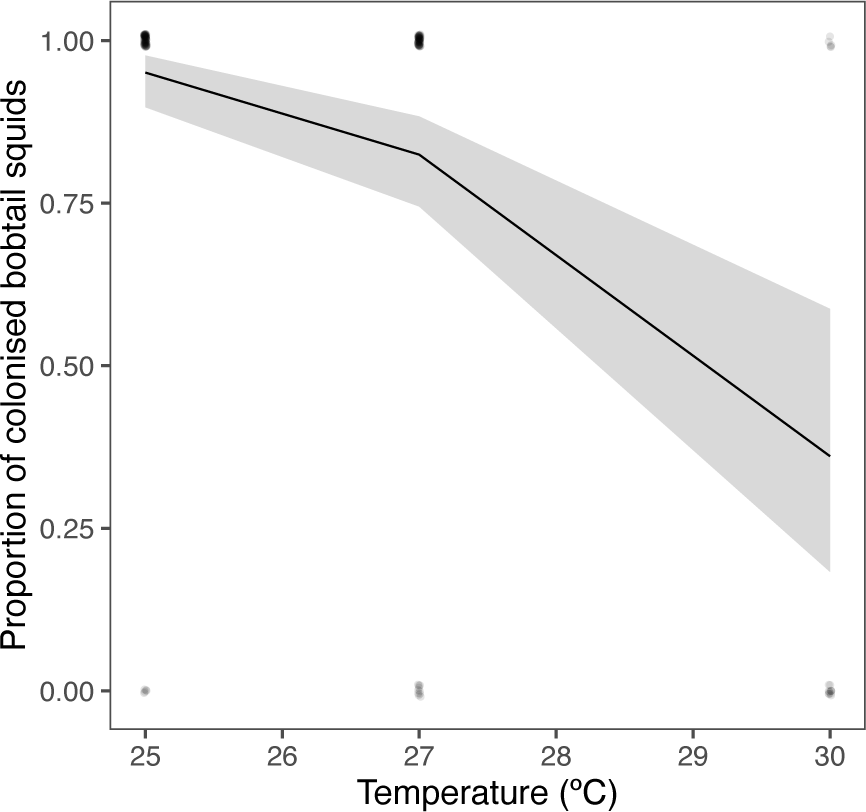
– Maintenance of the symbiosis at 48 h post-inoculation. The maintenance of the symbiont was analysed through a generalised mixed model from the binomial family as a function of the temperature. Line and shaded area are the back-transformed predicted means ± 95 % confidence interval from the model. The points are the raw data; ‘jitter’ was used to reduce overlap but should be considered 0 (non-colonised) or 100 (colonised).

### 3.3 Formation status of crypt 3

The presence of crypt 3 in the light organ (i.e., a formed crypt 3) depended on the temperature but not the developmental time (Supplementary Table 10.A). We considered crypt 3 as formed when the crypt space was present in at least one side of the light organ (Supplementary Figure 7). We assumed the absence of crypt 3 when the formation was limited to the beginning of the invagination (i.e., presence of the duct only; Figure 6.A). We found formed crypt 3 in all animals reared at 25 °C (Figure 6.B). However, the formation of this specific crypt significantly decreased with warming (p-value < 0.001; Supplementary Table 10.A-B), reaching 94.9 % at 27 °C (n_crypt3_27_/n_total_27_ = 37/39), to only 46.5 % at 30 °C (n_crypt3_30_/n_total_30_ = 20/43, Figure 6.B).

**Figure 6.**
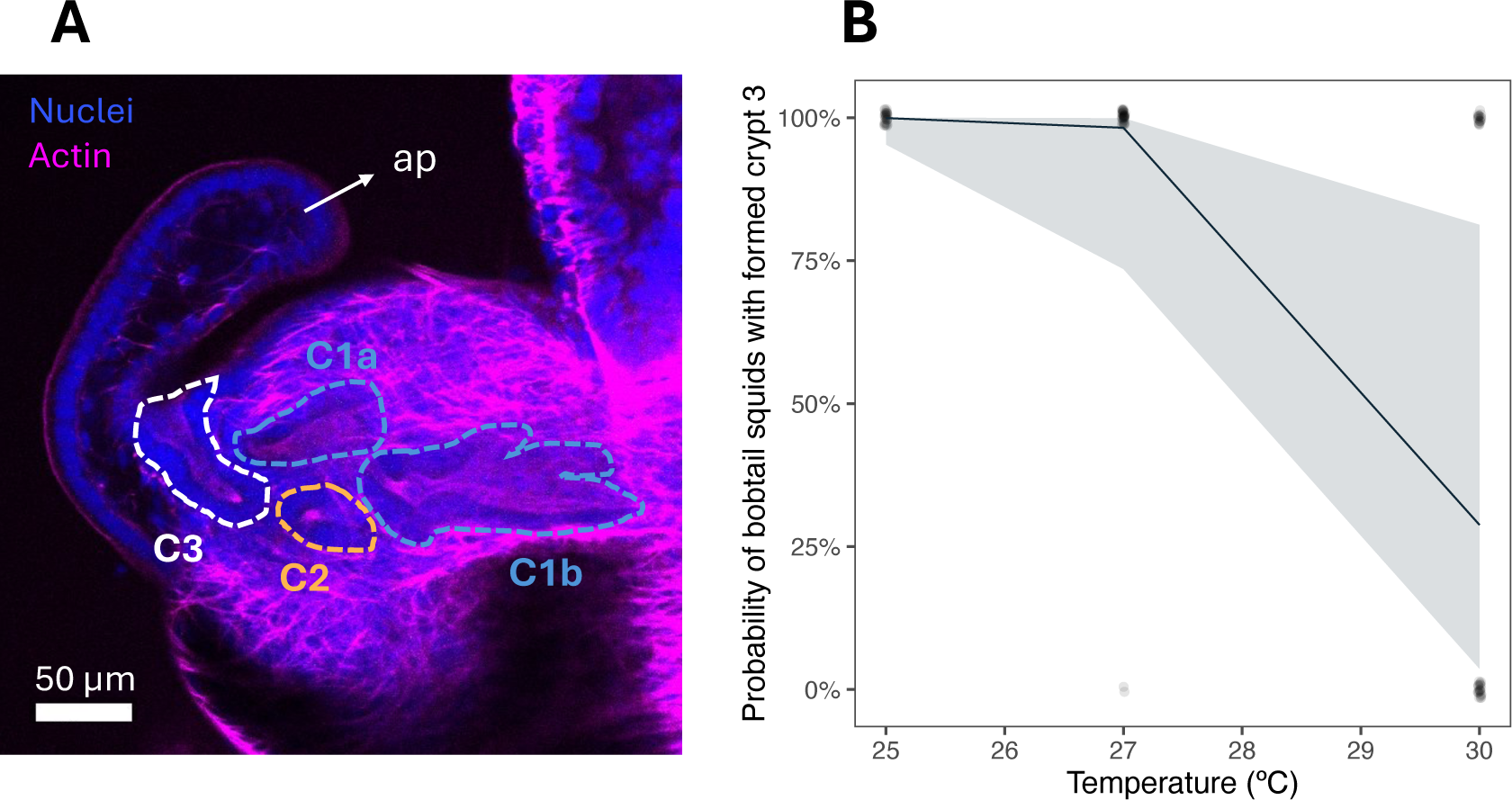
Formation of crypt 3 according to temperature. (A) Representative confocal image of a light organ with no crypt 3 present. The dotted lines delimit the three crypts: light blue = crypt 1 antechamber (C1a) and crypt 1 space (C1b), orange = crypt 2 space (C2) and white = crypt 3 invagination (C3). Nuclei are shown in blue, and F-actin in magenta. (B) Generalised mixed model from the binomial family for C3 formation as a function of the temperature. Line and shaded area are back-transformed predicted means ± 95 % confidence interval from the model. The points are the raw data; ‘jitter’ was used to reduce overlap but should be considered 0 (crypt 3 space not observed) or 100 (crypt 3 space observed).

## 4 Discussion

The present study revealed a significant negative effect of environmental heat stress on the establishment of the symbiosis during early ontogeny of the Hawaiian bobtail squid, *Euprymna scolopes*, with the luminescent bacterium *Vibrio fischeri*. Here, we showed that embryogenic exposure to increased temperature led to a significant decrease in the bobtail squid’s hatching success and development time, as described in squids and octopuses (Rosa et al., 2012, 2014; Repolho et al., 2014). Bobtail squid specimens reared under increased temperature (i.e., summer temperature and category IV MHW) tended to hatch earlier, probably linked to increased energy expenditure rates (Pimentel et al., 2012; Rosa et al., 2012).

In the late embryogenetic stages, the Hawaiian bobtail squid possesses an internal yolk sac that will be depleted in the first few days post-hatching (Lee et al., 2009; Moriano-Gutierrez et al., 2020). The reduced survival of newborns during the first 48 h post-hatching at 30 °C may be linked to a faster depletion of the internal yolk sac. In contrast, we did not observe a significant difference in survival at 27 °C compared to animals kept at 25 °C, which may be explained by the fact that 27 °C corresponds to the average temperature during the summer months, i.e., it is within the optimum thermal range of this bobtail squid species (Pörtner et al., 2017).

When monitoring the colonisation of *E. scolopes* by *V. fischeri*, we show that the initiation of this symbiosis is altered under warming conditions. Adaptation to stressful temperature by *Vibrio fischeri* was suggested to facilitate the colonisation of *E. scolopes* by this bacterium (Cohen et al., 2019). However, our results show that when the bobtail squid is itself exposed to the increased temperature during embryogenesis, there is a negative impact on its colonisation. This may be explained by the negative correlation between the expression of genes involved in the initiation of the symbiosis and increasing temperature (Otjacques et al., unpublished data). Therefore, the initiation of this squid-vibrio symbiosis may be negatively impacted under increased sea surface temperatures. Moreover, we observed a reduced luminescence in bobtail squids exposed to increased temperatures. This may be linked to an alteration in the production of autoinducer (AI). Indeed, the intensity of bioluminescence expressed by *V. fischeri* depends on the amount of AI produced to subsequently reach quorum sensing (Nealson, 1977; Boettcher & Ruby, 1995). However, the autoinducer-2 activity, used in bioluminescence regulation by some *Vibrio* species, was shown to be negatively impacted by increasing temperature (Zhang et al., 2008). In addition, in *V. fischeri*, an *N*-Acyl homoserine lactone (AHL) is another known AI involved in light production, for which the synthesis depends on the expression of the *luxI* gene (Boettcher & Ruby, 1990). Increasing temperatures also negatively affect the production of AHL (Hansen et al., 2015). Therefore, our results suggest that elevated temperatures may not only negatively impact the colonisation of the bobtail squid but also reduce the bioluminescent capacity of this organism.

Once colonised, the bobtail squids undergo a profound morphogenesis of the light organ, with the regression of the appendages (Montgomery & McFall-Ngai, 1994). The complete regression follows an irreversible signal that occurs when *V. fischeri* cells settle into the light organ’s crypt spaces (Doino & McFall-Ngai, 1995). Our results show that the irreversible signal exhibits the same regression rate across all treatments. This implies that the morphogenesis following colonisation does not depend on temperature. However, partial regression can be induced through the recognition of bacterial components, such as lipopolysaccharide (LPS; Foster et al., 2000; Koropatnick et al., 2004). Since we have performed the colonisation with natural (offshore) seawater, we can assume that the slight morphogenesis observed in non-colonised animals is due to the presence of LPS and other bacterial components in the surrounding seawater. Therefore, we suggest that temperature can indeed influence the regression of the light organ’s appendages, but only until the irreversible signal is reached.

In the squid–vibrio symbiosis, it is common for the bobtail squid to partially expel the symbiont into the environment. In fact, following the daily dawn light cue, the bobtail squids expel around 90% of the bacteria present in the crypt space, a process called ‘venting’ (Essock-Burns et al., 2020; Nyholm & McFall-Ngai, 2021). Moreover, when exposed to antibiotics, *E. scolopes* is also expected to vent, with the exception of the cells present in the crypt 3 (Essock-Burns et al., 2023). However, whereas the animal usually partially retains the symbiont, here we demonstrate that increased temperature provokes a complete curation of the bacteria from the light organ. In other words, bobtail squids exposed to higher temperatures do not maintain the symbiosis. Marine heatwaves have been shown to drive symbiont expulsion among symbiotic organisms. Therefore, our results are consistent with this phenomenon observed in corals (Lesser, 2011; Smith et al., 2023) or in sponges, and octocorals, where the expulsion of certain strains leads to a reorganisation of their microbiome (Prioux et al., 2023; Bell et al., 2023, 2024). Another explanation may come from the feeding status of the bobtail squids. Due to the presence of the internal yolk sac, we have kept our animals unfed. However, since we estimated the yolk sac to deplete faster under increased temperatures, we suggest that the animals may expel the bacteria to conserve their food reserves. This is consistent with the lack of symbiosis maintenance observed when bobtail squids were colonised with a mutant, which also led to a faster depletion of the yolk sac (Moriano-Gutierrez et al., 2020). Further investigation should be made to test this hypothesis, as we suggest that the symbiosis may be maintained depending on the ability of the animal to feed while the yolk sac is depleting.

Finally, our results demonstrate that embryos developing under a category IV MHW are significantly less likely to form the C3. A total of 30 stages are used to describe the embryogenesis of the Hawaiian bobtail squid, with crypt spaces beginning their formation sequentially merely a few days before hatching (Montgomery & McFall-Ngai, 1993; Lee et al., 2009). Moreover, there is a linear relation between developmental time and temperature, for which it is expected to reach certain embryonic stages sooner under increased temperatures, up to the final hatching stage (Jiang et al., 2020; Márquez et al., 2021, 2023). Here, the exposure to a category IV MHW may have resulted in premature hatching at the embryonic stage A29, when the third pair of crypts begins its formation (Montgomery & McFall-Ngai, 1993). Although the animal can hatch healthily at that stage (Lee et al., 2009), it is admitted that the C3 provides resilience to the bobtail squid early stages under stress by providing a reserve of bacteria ready to recolonise the animal when conditions improve (Essock-Burns et al., 2023). However, since we identified a loss of symbiosis maintenance with increased temperatures, we suspect that the absence of the C3 may indeed impair the resilience of *E. scolopes* to maintain a functional symbiosis.

In summary, our results confirm that the Hawaiian bobtail squid *Euprymna scolopes* is a species sensitive to increased temperatures during early ontogney, resulting in decreased hatching success and increased mortality following hatching when exposed to extreme temperatures (e.g., category IV marine heatwave). We also indicate that the symbiosis between *E. scolopes* and the bacterium *Vibrio fischeri* may be in jeopardy under future climate conditions and the increase in marine heatwave events. We demonstrate that increased temperatures are detrimental to the initiation of this symbiosis and lead to a reduced likelihood of maintaining the symbiont. Moreover, the loss of the crypt space 3 may diminish the symbiotic resilience of this species to external perturbation. In addition to the impact on corals and other marine organisms, our research suggests that temperature can also alter the initiation and establishment of bacterial symbiosis in a cephalopod. Consequently, this could negatively affect the behaviour of this species because of the loss of its ability to camouflage at night by counterillumination. Our findings highlight the need for more research on microbial symbiosis when assessing the impact of climate change on marine holobionts.

## 5 Ethics statement

This experiment was performed in compliance with the ethical standards of the Institutional Animal Care and Use Committee and the approved protocol IA22-1842 at the California Institute of Technology.

## 6 Data availability

All datasets and R codes will be available in the Figshare repository upon publication.

## 7 Author contributions

Conceptualisation: E.O., R.R., M.M-N.; Data curation: E.O.; Formal analysis: E.O., T.M.; Funding acquisition: E.R., M.M-N.; Investigation: E.O., B.J.; Methodology: E.O., E.R., M.M-N., R.R.; Project administration: E.R., M.M-N.; Resources: E.R., M.M-N.; Supervision: E.R., J.X., R.R., M.M-N.; Visualization: E.O.; Writing – original draft: E.O., M.M-N., R.R.; Writing – review & editing: E.O., B.J., T.M. J.X., E.R., R.R., M. M-N.

All authors gave final approval for the submission of this manuscript

## 10 Acknowledgments

The authors would like to thank all the lab members from the McFall-Ngai and Ruby labs for their help and great discussions over the results of this manuscript.

