## Supplementary Figures for "Climate-driven warming disrupts the symbiosis of bobtail squid *Euprymna scolopes* and the luminous bacterium *Vibrio fischeri*"

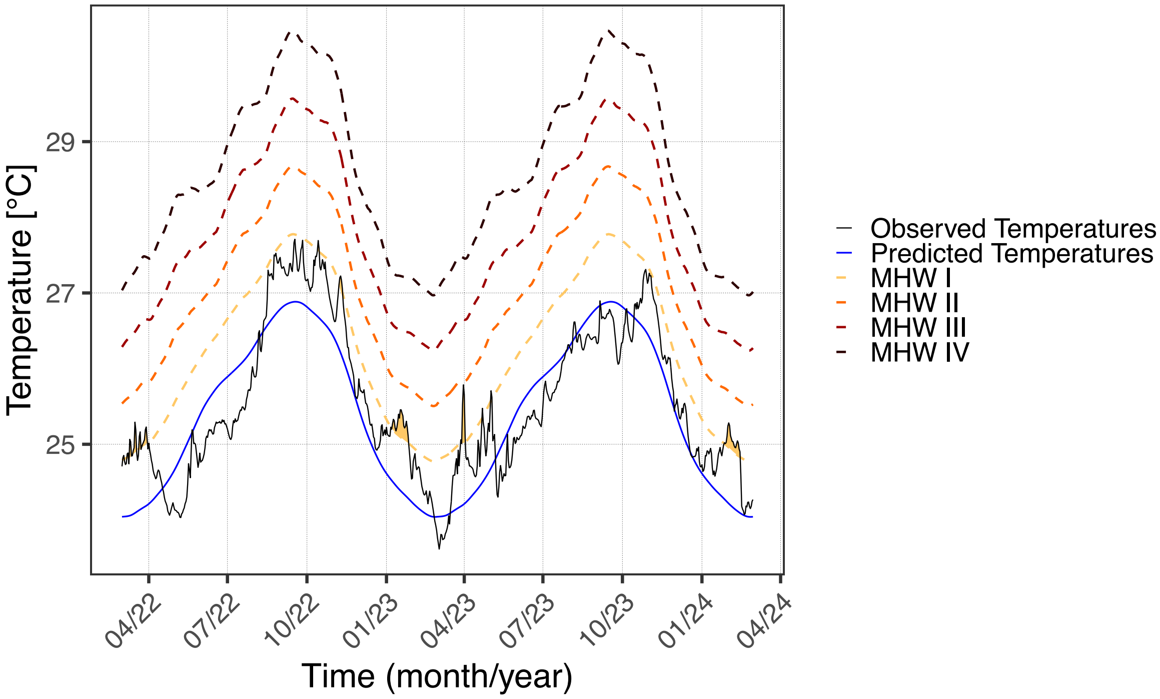


**Supplementary Figure 1** – Climatology for 03/2022-02/2024. Sea Surface Temperatures (SSTs) collected from the Copernicus Marine Service (21.25 ºN, 157.75 ºW) between 31/01/1993 and 29/02/2024. Daily average measured SSTs is in black (Observed temperatures). The climatology line (blue) represents the expected SSTs, modelled using the 31-year dataset. The coloured dashed lines are the expected SSTs during different marine heatwaves (MHW), ranging from category I (MHW I) to category IV (MHW IV). The coloured areas represent observed marine heatwaves, when the measured SSTs are above the threshold for longer than 5 consecutive days.

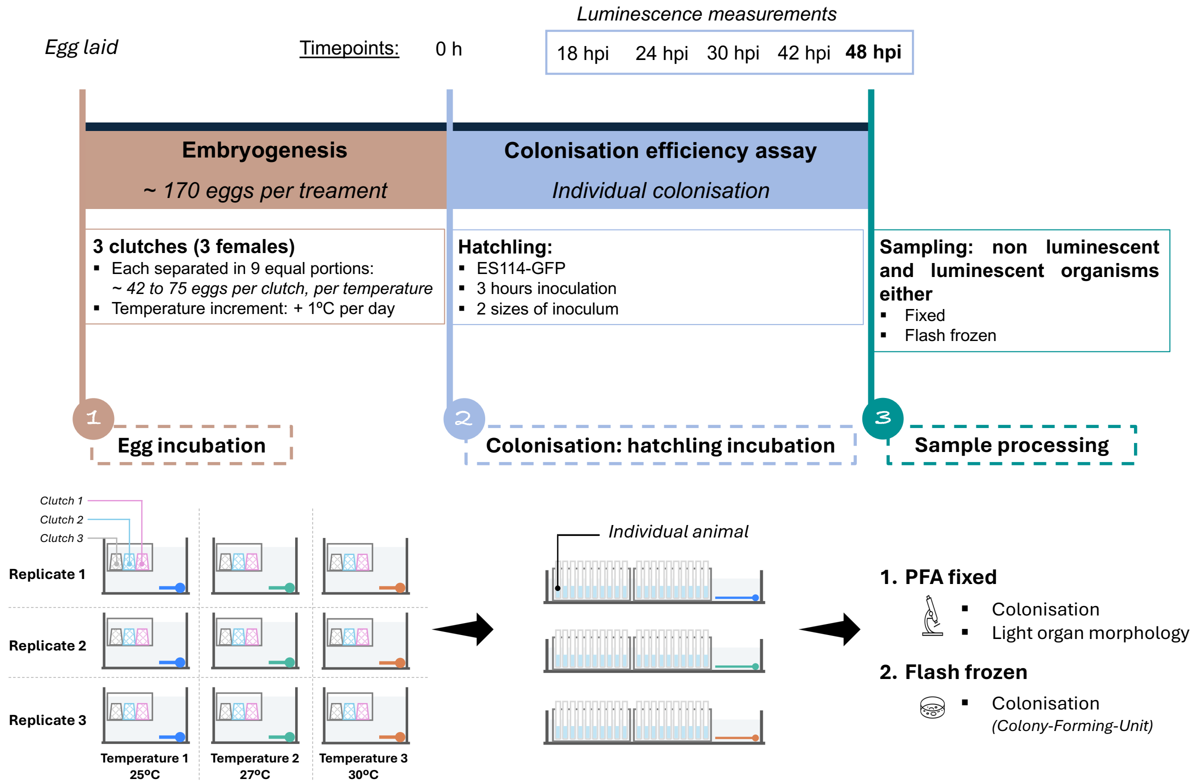


**Supplementary Figure 2** – Experimental design. (1) The **egg incubation** started on the day the clutches were laid. The temperature increment followed a 1ºC increase per day, until reaching the final temperature corresponding to each treatment. (2) The **colonisation** of the animals was performed individually, and the temperature was kept constant, according to the rearing temperature during embryogenesis. (3) At 48 hours post-inoculation (hpi), luminescent and non-luminescent were either fixed in 4% paraformaldehyde or flash frozen. The **sample processing** was followed according to the sampling method.

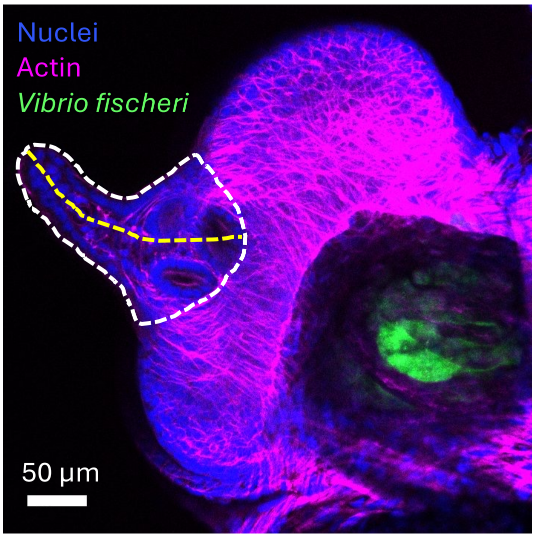


**Supplementary Figure 3** – Confocal image of a light organ from a colonised squid. Representative image of the measurement for the appendage length (yellow dotted line) and the appendage area (white dotted line).

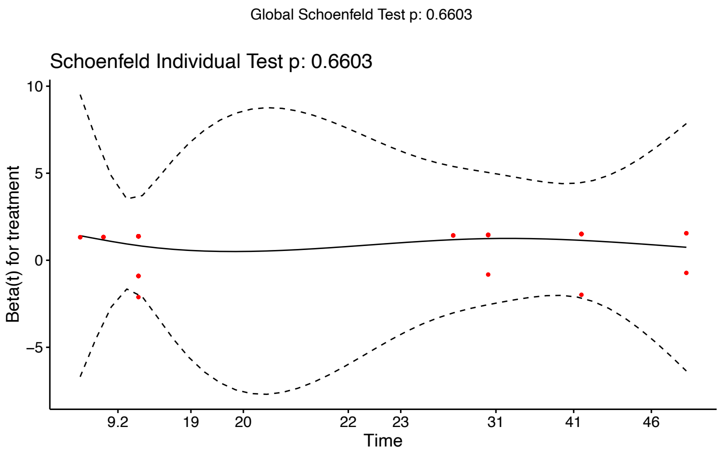


**Supplementary Figure 4 –** Smoothed spline plots of Schoenfeld residuals of the survival Cox mixed effects model relative to time.

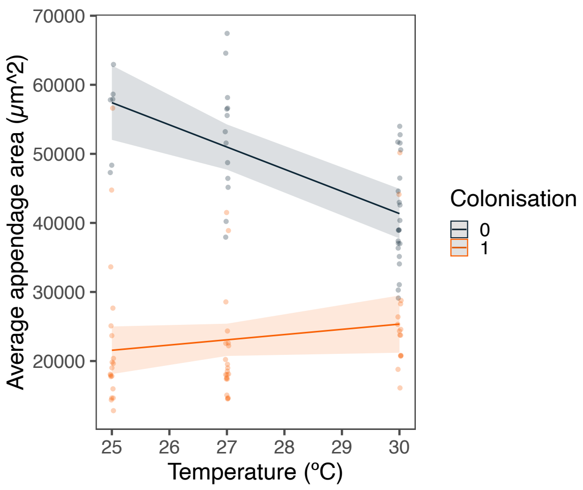


**Supplementary Figure 5 –** Light organ's appendages area according to temperature and colonisation. The morphology of the light organ, the area of the appendages more specifically, was analysed through a generalised mixed model from the Gaussian family, in function of the temperature and the colonisation. Non-colonised (0) bobtail squids are in black and colonised (1) bobtail squids in red. Back-transformed predicted means ± 95% confidence interval from the model and raw data values are presented. The areas of both appendages were averaged for each individual.

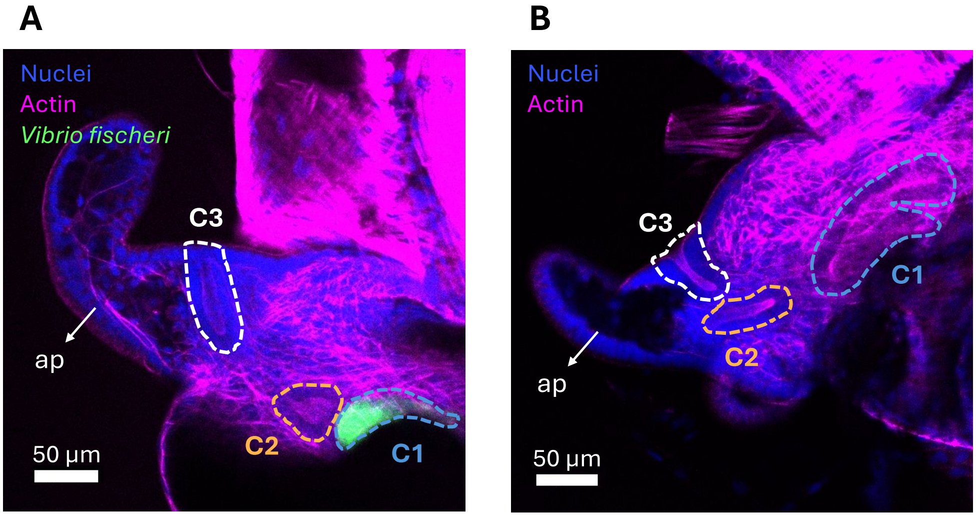


**Supplementary Figure 6** – Confocal images of a light organ, representative for the maintenance (A) or the non-maintenance (B) of the symbiosis (regression of the arm is visible but bacteria are absent from all crypts). The crypt spaces are delimited by the dotted lines (crypt 1 = C1 is in white, crypt 2 = C2 is in orange, and crypt 3 = C3 is in light blue). The arrow shows the regressed appendage (ap). The nuclei are shown in blue, F-actin in magenta, and the bacterium *Vibrio fischeri* is represented in green.

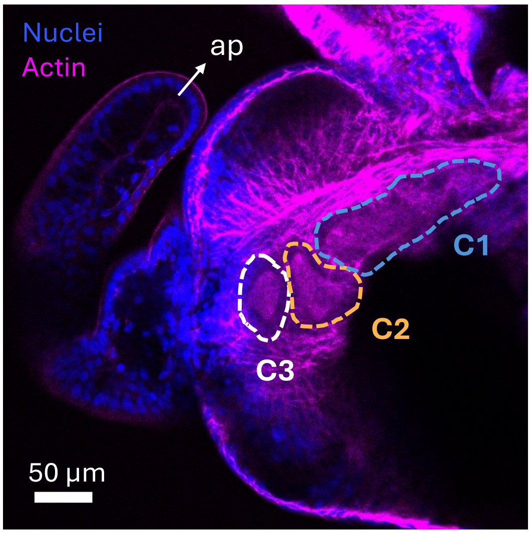


**Supplementary Figure 7** – Confocal image of a light organ, representative image for the formation of the three crypt spaces. The three crypts are delimited by the dotted lines (crypt 1 = C1 is in white, crypt 2 = C2 is in orange and crypt 3 = C3 is in light blue). The nuclei are shown in blue and F-actin in magenta.
